# Therapeutic Knock-in Genome Editing Using Single AAV Vectors in Mouse Models of Inherited Liver Disease

**DOI:** 10.1101/2025.07.30.667771

**Authors:** Khishigjargal Batjargal, Tomoki Togashi, Yuji Kashiwakura, Nemekhbayar Baatartsogt, Kosuke Tsuchida, Takahiro Sato, Morisada Hayakawa, Kiwako Tsukida, Kazuhiro Muramatsu, Atsushi Hoshino, Osamu Nureki, Tsukasa Ohmori

**Affiliations:** Department of Biochemistry, Jichi Medical University School of Medicine, 3311-1 Yakushiji, Shimotsuke, Tochigi 329-0498, Japan; Center for Gene Therapy Research, Jichi Medical University, 3311-1 Yakushiji, Shimotsuke, Tochigi 329-0498, Japan; Department of Surgery, Jichi Medical University School of Medicine, 3311-1 Yakushiji, Shimotsuke, Tochigi 329-0498, Japan; Department of Pediatrics, Jichi Medical University School of Medicine, 3311-1 Yakushiji, Shimotsuke, Tochigi 329-0498, Japan; Department of Cardiovascular Medicine, Graduate School of Medical Science, Kyoto Prefectural University of Medicine, Kyoto 602-8566, Japan; Department of Biological Sciences, Graduate School of Science, The University of Tokyo, 7-3-1 Hongo, Bunkyo-ku, Tokyo 113-0033, Japan

**Author notes:** **Corresponding Author:** Tsukasa Ohmori, M.D., Ph.D., Department of Biochemistry, Jichi Medical University School of Medicine, 3311-1 Yakushiji, Shimotsuke, Tochigi 329-0498, Japan. These authors contributed equally (Khishigjargal Batjargal, Tomoki Togashi).

## Abstract

Gene knock-in therapy has the potential to cure inherited liver diseases but is limited by low efficiency and delivery complexity. Here, we developed a single adeno-associated virus (AAV) vector system comprising a compact CRISPR effector, enAsCas12f, a guide RNA, and a donor template to enable therapeutic genome editing via non-homologous end joining (NHEJ). We targeted the system to the murine *Alb* locus and applied it to mouse models of hemophilia B, protein C (PC) deficiency, and ornithine transcarbamylase (OTC) deficiency. NHEJ-mediated knock-in showed higher efficiency than homology-directed repair, with successful therapeutic gene insertion in both neonatal and adult mice. The strategy restored plasma factor IX activity in hemophilia B (*F9^−/−^*) mice, prolonged survival of PC-deficient (*Proc^−/−^*) mice, and prevented hyperammonemia and weight loss in OTC-deficient (*Otc^spf-ash^*) mice upon high protein challenge. Importantly, gene integration was restricted to the liver, with no evidence of germline transmission. This compact, all-in-one AAV knock-in platform simplifies vector production, enables efficient delivery, and achieves reliable transgene expression *in vivo*. Our findings highlight the potential of liver-targeted knock-in genome editing as a transplant-independent treatment for neonatal-onset metabolic diseases, offering a clinically feasible path towards curative gene therapies for a wide range of monogenic liver disorders.

## Introduction

A number of essential physiological functions occur in the liver, including metabolism, detoxification, and the synthesis of plasma proteins. Inherited abnormalities of these functions can lead to life-threatening conditions in the neonatal period (1). For example, inherited deficiency of protein C (PC), an anticoagulant protein produced in the liver, predisposes systemic fatal thrombosis (2), while inherited deficiency of ornithine transcarbamylase (OTC), a key enzyme of the urea cycle, causes hyperammonemia, leading to diverse and severe neurological disturbances (3, 4). Conservative medical management of these conditions usually has limited therapeutic efficacy and liver transplantation, including living donor transplantation, is currently the only definitive curative option for severe cases (5). However, liver transplantation is associated with significant limitations, including a critical shortage of donors, requirement for lifelong immunosuppression (6), and substantial perioperative risks (7). Living donor liver transplantation may become an alternative option, but it imposes considerable physical and psychological stress on the donor (8). These challenges underscore the urgent need for curative therapies in the neonatal period that do not rely on transplantation.

Gene therapy has emerged as a promising modality for curing congenital disorders. Indeed, adeno-associated virus (AAV) vector-mediated gene therapy has shown promise in treating several monogenic liver disorders and is currently under clinical investigation for several diseases, including OTC deficiency and hemophilia (9, 10). AAV vectors efficiently deliver exogenous functional genes to hepatocytes by intravenous administration, resulting in the expression of therapeutic proteins in the liver. However, the AAV genomes within the cells persist primarily as episomal DNA and are rarely integrated into the host genome (11). These AAV vector characteristics limit the duration of exogenous gene expression in the liver because hepatocyte proliferation gradually dilutes the AAV genome over time (12). Therefore, AAV-mediated gene therapy does not currently offer a curative solution for severe neonatal-onset metabolic liver diseases.

Genome editing technologies that directly target chromosomal DNA offer a potential breakthrough to overcome the limitations of AAV vectors for the treatment of severe neonatal-onset metabolic liver diseases. Knock-in genome editing enables the integration of therapeutic genes into the host genome, ensuring stable and long-term gene expression, even in proliferating hepatocytes. Indeed, knock-in genome editing can ameliorate disease phenotypes in various models of liver disorders (13–15). Efficient knock-in of therapeutic genes typically requires the induction of double-strand breaks by nucleases, such as Cas9, which activates the DNA repair machinery of the cell. Current knock-in approaches require two separate components—one for the nuclease and one for the donor DNA—making clinical translation difficult because of delivery inefficiencies and manufacturing complexity.

To address these limitations, we previously developed an engineered variant of AsCas12f (enAsCas12f-HKRA), which is less than one-third of the size of SpCas9 but exhibits comparable editing activity (16). Using this compact editor, we demonstrated that homology-directed repair (HDR)-mediated knock-in can be achieved with a single AAV vector, successfully restoring factor IX (FIX) expression in a hemophilia B mouse model *in vivo* (16). However, the efficiency of HDR-mediated knock-in remains low (17), and further optimization is needed to achieve effective therapeutic outcomes. In this study, we enhanced the efficiency of knock-in genome editing using AsCas12f and evaluated its therapeutic efficacy in murine models of hemophilia B, PC deficiency, and OTC deficiency. This genome editing platform holds promise as a clinically feasible and transplant-independent strategy for treating a broad range of neonatal-onset monogenic liver diseases.

## Results

### Knock-in strategy with a single AAV vector targeting the Alb locus in hemophilia B mice

We previously demonstrated that therapeutic genes can be inserted into the *Alb* locus in mouse liver cells via HDR using a single AAV vector containing enAsCas12f-HKRA and a donor sequence (16). However, the *in vivo* knock-in efficiency via HDR is lower compared with that via non-homologous end joining (NHEJ) (18–20). To compare the knock-in efficiency mediated by HDR and NHEJ in AsCas12f-mediated knock-in strategies, we first selected single guide RNAs (sgRNAs) to induce double-strand breaks at the respective loci for each knock-in (sgRNA1 [intron 14] for HDR and sgRNA5 [intron 13] for NHEJ) (Supplemental Figure 1). We designed a donor sequence to insert a human *F9* (*hF9 [R338L])* cDNA, a gain-of-function variant of the *F9* gene, at the double-strand break sites. The R338L variant, which has been used in gene therapy for hemophilia B, displays approximately eightfold higher specific activity than the wild-type protein (21). Homologous arms (500 bp) were appended to both ends of the cDNA for in-frame knock-in of *hF9 (R338L)* via HDR targeting *Alb* intron 14, but not via NHEJ targeting intron 13 (Figure 1A, B).

**Figure 1.**
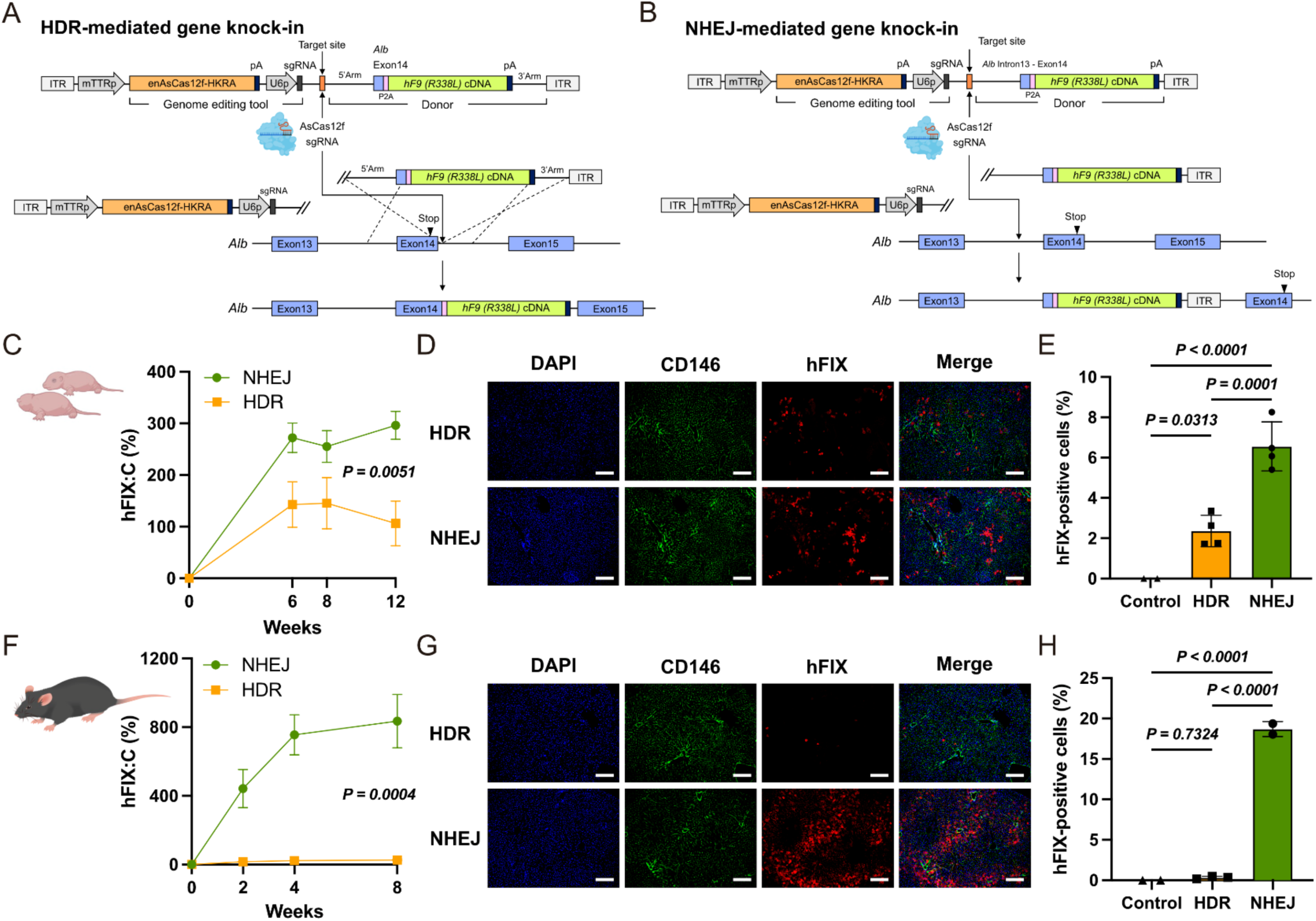
Knock-in genome editing therapy using a single AAV vector for hemophilia B neonatal and adult mice. (**A, B**) Schematic representation of knock-in genome editing using a single adeno-associated virus (AAV) vector. The single AAV vector was composed of a genome editing cassette comprising enAsCas12f-HKRA driven by the mouse *Ttr* promoter and a single guide RNA (sgRNA) targeting *Alb* intron 14 or 13 driven by the U6 promoter, and a donor template containing *Alb* and a human *F9 (R338L)* cDNA. An sgRNA target sequence was inserted upstream of the donor template to enable cleavage by enAsCas12f-HKRA. The stop codon of *Alb* included in the donor template was replaced with a P2A sequence to enable separate translation of albumin and hFIX. (**A**) Knock-in of *hF9 (R338L)* cDNA via homology-directed repair (HDR) into *Alb* intron 14 using double-strand breaks. (**B**) Knock-in of *hF9 (R338L)* cDNA into *Alb* intron 13 via non-homologous end joining (NHEJ). (**C–H**) A single AAV vector (3 × 10¹¹ vg/neonatal mouse, 1 × 10¹^2^ vg/adult mouse) was administered to neonatal and adult hemophilia B mice. (**C, F**) Plasma hFIX activity (hFIX:C). Values represent the mean ± SD (n = 4). *P* values were determined using two-way ANOVA. (**D, G**) Immunofluorescence images of hFIX in the liver were photographed using a BZ-X700 fluorescence microscope (Keyence). Scale bar: 200 µm. (**E, H**) Quantification of FIX-positive cells. Values represent the mean ± SD (n = 4, except negative control [n = 2], in neonatal experiments; n = 3, except control and NHEJ [n = 2], in adult experiments). *P* values were determined using one-way ANOVA with Tukey’s multiple comparisons test. ITR, inverted terminal repeat.

We constructed two AAV8 vectors harboring enAsCas12f-HKRA and a donor sequence: one targeting *Alb* intron 14 for HDR-mediated donor insertion, the other targeting *Alb* intron 13 for NHEJ-mediated donor insertion. These AAV vectors were intraperitoneally injected into neonatal FIX-deficient hemophilia B mice (Figure 1A). The increase in plasma hFIX levels was significantly higher in mice treated with the NHEJ-mediated knock-in, compared with those treated with HDR-mediated knock-in (Figure 1B). To quantify the knock-in efficiency, we assessed hFIX-positive cells in the liver by immunostaining (Figure 1D). NHEJ-mediated knock-in resulted in more hFIX-positive cells in the liver than HDR-mediated knock-in (Figure 1D, E). These results indicate that NHEJ-mediated knock-in targeting intron 13 is more efficient than HDR-mediated knock-in for achieving genome-editing therapy in neonatal mice. Next, we compared the knock-in strategies via HDR and NHEJ in adult hemophilia B mice. An increase in plasma hFIX activity (hFIX:C) was observed only in the NHEJ-mediated knock-in group (Figure 1F) and the number of hFIX-positive cells in NHEJ knock-in mice was 18.71% (Figure 1G, H). Plasma albumin levels were not affected by the knock-in genome editing (Supplemental Figure 2).

We assessed the knock-in pattern of the donor sequence in the genomic DNA of the neonatal mice by PCR. We were able to distinguish between HDR- and NHEJ-mediated insertions based on the size of the PCR products. When targeting intron 14, HDR resulted in a 1.5 kbp product, whereas NHEJ yielded a 2.2 kbp product. In contrast, when targeting intron 13, NHEJ produced a 1.5 kbp PCR product (Supplemental Figure 3A). The insertion of the *hF9 (R338L)* cDNA into the target region was confirmed by PCR amplification (Supplemental Figure 3). When targeting intron 14, both HDR- and NHEJ-mediated knock-in events were detected by PCR, with NHEJ being more prominent (Supplemental Figure 3B). Targeting intron 13 only resulted in the amplification of the PCR product corresponding to NHEJ-mediated integration (Supplemental Figure 3B).

### Genome editing prolongs the survival of protein C-deficient mice

We next evaluated our strategy for fatal monogenic disorders that have the potential to be treated by targeting the liver. First, we applied our approach to PC deficiency. PC is an anticoagulant protein produced in the liver that inactivates activated coagulation factor V (FV) and factor VIII (FVIII), thereby inhibiting thrombus formation (22). Abnormalities in both alleles of *PROC*, the gene that encodes PC, cause neonatal purpura fulminans, a life-threatening condition characterized by systemic thrombosis shortly after birth (23). We previously achieved neonatal genome editing using a two-AAV vector system, with one AAV vector carrying Cas9 and the other containing a donor sequence to express an engineered active form of mouse PC (mPC-2RKR) (24). Here, we used a strategy in which AsCas12f and the donor sequence were incorporated into a single AAV vector. We then examined the effect of this knock-in genome editing therapy on PC-deficient (*Proc^−/−^*) mice.

We first evaluated whether engineered human PC (hPC-2RKR) is a suitable donor sequence for genome editing to rescue the phenotype of *Proc^−/−^*mice. We expressed wild-type hPC or hPC-2RKR in wild-type mice using an AAV vector and evaluated their effects on anticoagulant function and other related parameters (Supplemental Figure 4). AAV vectors expressing either wild-type hPC or hPC-2RKR were intravenously administered to adult wild-type mice at varying doses. In mice expressing wild-type hPC, both hPC activity (hPC:C) and antigen levels (hPC:Ag) increased in a dose-dependent manner, whereas hPC-2RKR showed minimal antigen increase and undetectable activity (Supplemental Figure 4A, B). This is likely because activated PC is rapidly inactivated by various plasma factors (24), resulting in an extremely short half-life (25). However, hPC-2RKR prolonged activated partial thromboplastin time and reduced FV activity (FV:C) in a dose-dependent manner but did not affect FVIII activity (FVIII:C), indicating stronger anticoagulant effects even in mice (Supplemental Figure 4C–E). A tail clip assay showed similar bleeding patterns for wild-type hPC and hPC-2RKR treated mice (Supplemental Figure 4F, G), and liver AAV vector copy numbers were comparable between groups (Supplemental Figure 4H).

To determine the vector dose for neonatal genome editing to treat *Proc^−/−^* mice, we intraperitoneally administered AAV8 vectors harboring enAsCas12f-HKRA and donor targeting *Alb* intron 13 for NHEJ-mediated *PROC* cDNA insertion to wild-type neonatal mice at three vector doses. Plasma hPC:C at 6 weeks after vector injection increased in a dose-dependent manner (Figure 2A). We selected a dose of 1.0 × 10^11^ vg/mouse because this was expected to provide a sufficient plasma hPC level for phenotype correction. In addition to the AAV vector for knock-in genome editing, we injected anti-FVIII antibodies into *Proc^+/−^* pregnant female mice and their neonates shortly after birth, as described previously (Figure 2B) (24). The survival rates of *Proc^−/−^* mice with hPC or hPC-2RKR knock-in neonatal genome editing were 33.3% and 57.1%, respectively (Figure 2C). As expected, hPC:C was elevated only in the hPC knock-in group, while it remained below the detection limit in the hPC-2RKR knock-in group (Figure 2D). In contrast, FV:C tended to decrease only in the hPC-2RKR knock-in group (Figure 2E), while FVIII:C did not change (Figure 2F). These results indicate that using the functionally engineered hPC as the donor sequence improved therapeutic outcomes.

**Figure 2.**
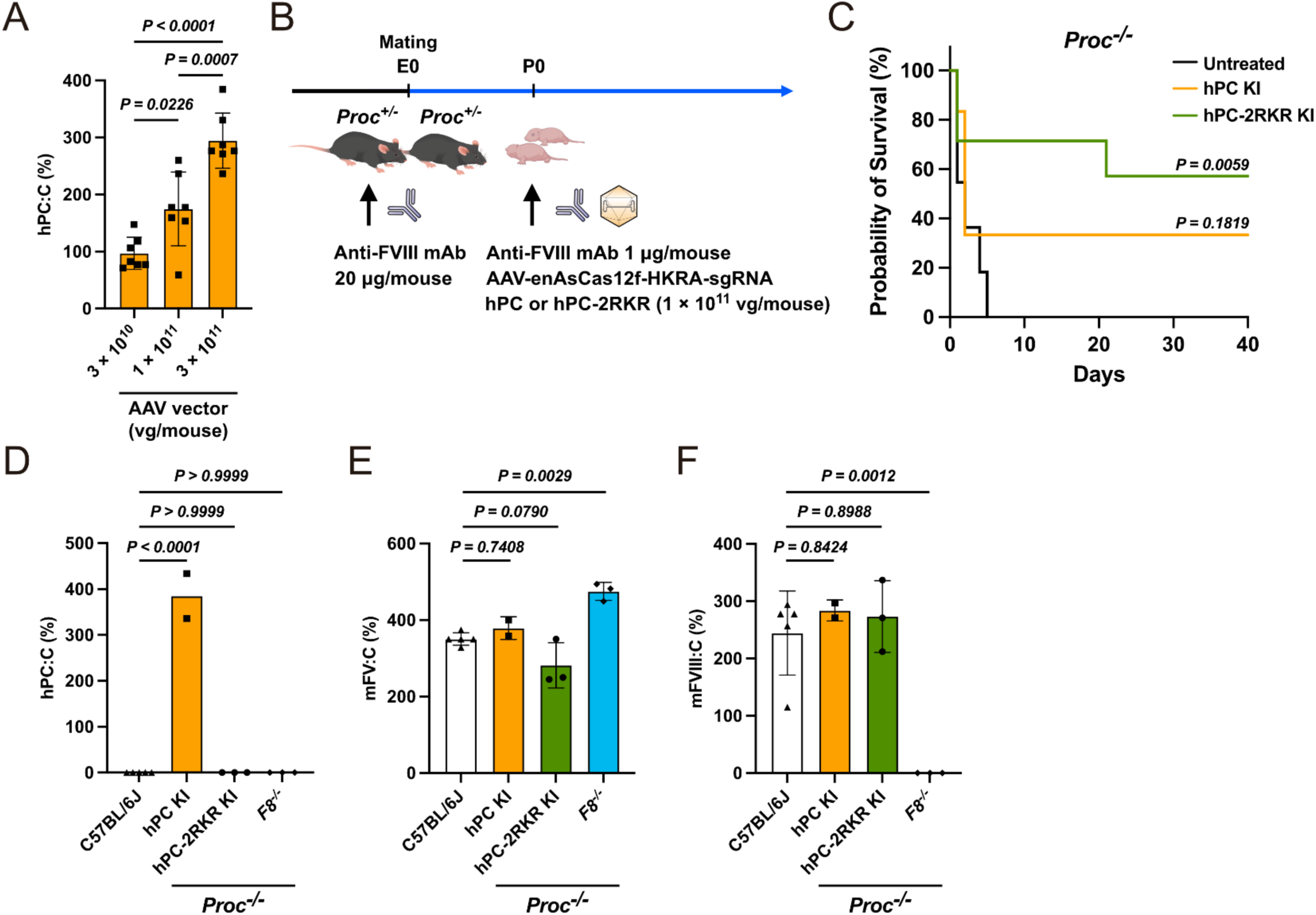
Rescue of the protein C-deficient mouse phenotype by knock-in genome editing with a single AAV vector. (**A**) Wild-type neonatal mice were intraperitoneally injected with varying doses of a single AAV vector containing enAsCas12f-HKRA-sgRNA-*PROC* cDNA. Human protein C (PC) activity (hPC:C) was measured 6 weeks after vector injection. Values represent the mean ± SD (n = 7). *P* values were determined using one-way ANOVA with Tukey’s multiple comparisons test. (**B**) *Proc^+/-^*female mice, mated with *Proc^+/−^* male mice, received an anti-factor VIII (FVIII) antibody every 7 days until delivery. The single AAV vector for knock-in hPC or hPC-2RKR (1.0 × 10^11^ vg/mouse) and the anti-FVIII antibody (1 µg/mouse) were injected into all neonatal mice. (**C**) Kaplan–Meier survival curves for *Proc^−/−^* mice (untreated, n = 13; hPC KI, n = 6; hPC-2RKR KI, n = 7). The survival curve for untreated *Proc^−/−^* mice was taken from a previous study (24). *P* values were determined using the log-rank test with comparison to untreated *Proc^−/−^* mice. (**D–F**) Plasma hPC:C (**D**), mouse factor V activity (mFV:C; **E**), and mouse factor VIII activity (mFVIII:C; **F**). Values represent the mean ± SD (n = 3–5, except hPC KI [n = 2]). *P* values were determined using one-way ANOVA with Tukey’s multiple comparisons test and comparison to C57BL/6J mice. KI, knock-in; vg, vector genome.

### Genome editing corrects the phenotype of Otc-deficient mice

The liver plays a crucial role in converting ammonia, a byproduct of amino acid catabolism, into urea through the urea cycle (also known as the ornithine cycle). While genetic abnormalities in several enzymes of the urea cycle can lead to hyperammonemia in newborns, OTC deficiency is the most prevalent (26). OTC deficiency is inherited in an X-linked recessive manner and is often associated with severe neonatal-onset phenotypes. Therefore, we investigated whether a knock-in strategy using a single AAV vector could improve the phenotype in OTC-deficiency model mice using adult *Otc^spf-ash^* mice, which have a spontaneous variant of the *Otc* gene (27). *Otc^spf-ash^* mice are viable and fertile, but exhibit hyperammonemia and weight loss when fed a high-protein diet (28, 29).

First, we generated three codon-optimized human *OTC* (*hOTC*) cDNAs, each conjugated with a FLAG tag sequence, and inserted them into the pcDNA3 plasmid. We then selected the sequence that exhibited the most efficient expression in human embryonic kidney (HEK) 293 cells (CO3 sequence in Supplemental Table 1). We injected a single AAV vector into adult male *Otc^spf-ash^* mice (3 × 10¹¹ or 1 × 10¹² vg/mouse) to insert the *hOTC* cDNA into the *Alb* intron 13 locus. Five weeks after AAV vector injection, the mice were challenged with a high-protein diet (Figure 3A). Untreated mice showed significant weight loss and a marked increase in blood ammonia levels (Figure 3B, C). However, administration of the AAV vector for genome editing predominantly prevented both weight loss and the elevation of blood ammonia (Figure 3B, C). hOTC enzyme activity in the liver was significantly increased at a titer of 1 × 10^12^ vg/mouse (Figure 3D). AAV-injected *Otc^spf-ash^* mice also had increased levels of hOTC protein and mRNA (Figure 3E–G). The knock-in efficiency assessed by immunofluorescence was estimated to be 11.75 ± 5.56% at a titer of 3 × 10¹¹ vg/mouse and 15.58 ± 3.90% at a titer of 1 × 10^12^ vg/mouse (Figure 3I).

**Figure 3.**
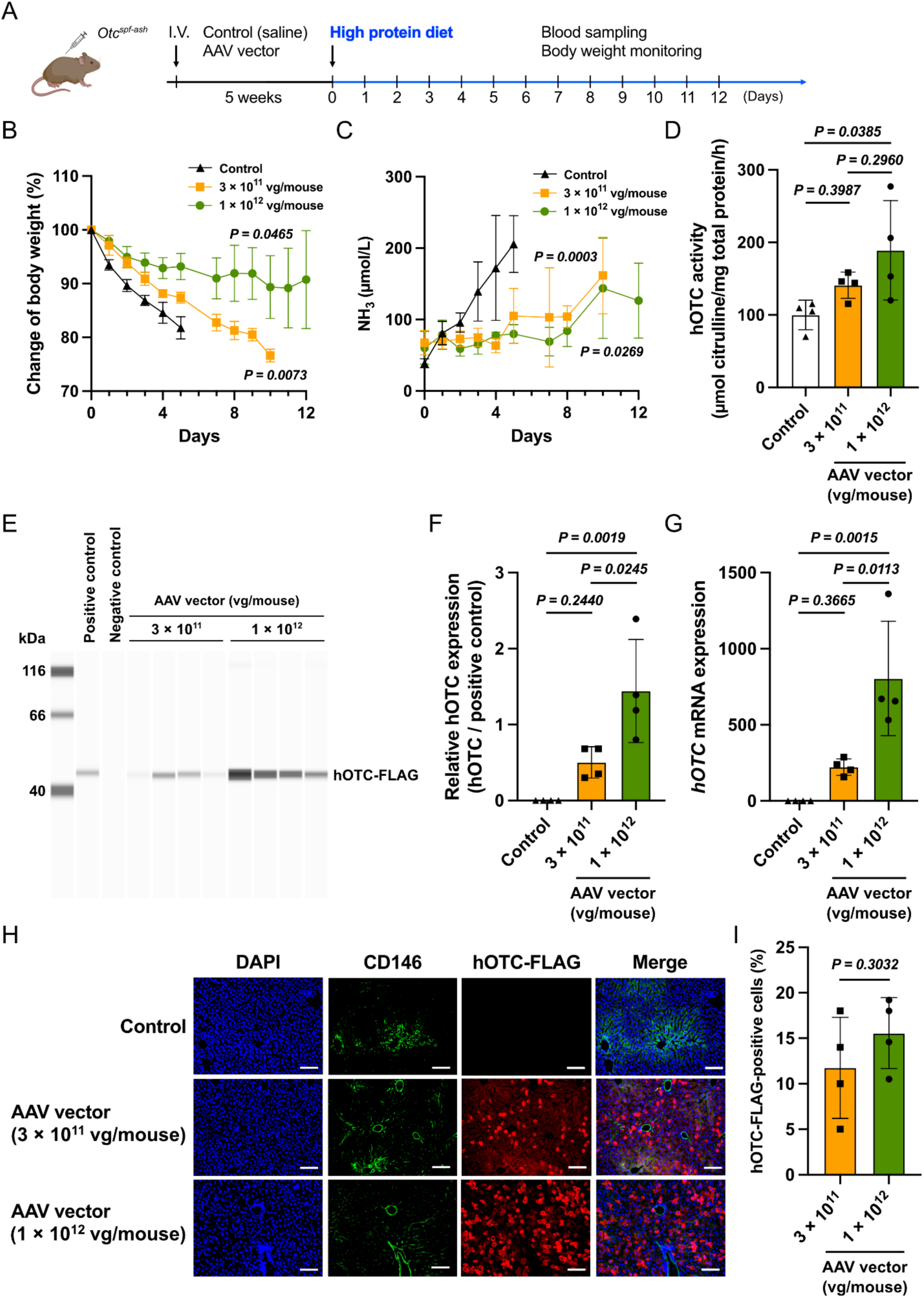
Rescue of the *Otc^spf-ash^* mouse phenotype by knock-in genome editing therapy. (**A**) Experimental design. Male *Otc^spf-ash^*mice were intravenously injected with AAV vector containing enAsCas12f-HKRA-sgRNA-*OTC* cDNA at 3 × 10^11^ or 1 × 10^12^ vg/mouse. Five weeks after vector injection, mice were challenged with a high protein diet (40% kcal). Body weight and blood ammonia (NH_3_) levels were determined at indicated time points for 12 days after the challenge. (**B, C**) Changes in body weight (% of that on day 0) (**B**) and blood NH_3_ concentrations (**C**) after the challenge. Values represent the mean ± SD (n = 4). The *P* values were determined by two-way ANOVA in comparison with the control. (**D**) hOTC enzyme activity in the liver of *Otc^spf-ash^*mice with or without (Control) knock-in genome editing. The *P* values were determined using one-way ANOVA with Tukey’s multiple comparisons test. (**E**) Immunoblotting analysis of liver hOTC protein in *Otc^spf-ash^* mice with or without (negative control) knock-in genome editing. Cell lysates of HEK293 cells transfected with pcDNA3 harboring hOTC were used as a positive control. (**F**) Quantification of hOTC protein. The *P* values were determined using Tukey’s multiple comparisons test. (**G**) *hOTC* mRNA levels in liver were analyzed by qPCR and normalized against *Gapdh* mRNA levels. Values are expressed as means ± SD. The *P* values were determined using one-way ANOVA with Tukey’s multiple comparisons tests. (**H**) Immunofluorescence images of hOTC in the liver were photographed using a BZ-X700 fluorescence microscope (Keyence). Scale bar: 100 µm. (**I**) Quantification of hOTC-positive cells in liver. The *P* value was determined using the two-tailed Student’s *t*-test. vg, vector genome.

To assess the correction of neonatal OTC deficiency using our strategy, we attempted to generate knockout model mice by deleting exon 2 of *Otc* through genome editing (Supplemental Figure S6A). However, we were unable to obtain any neonatal *Otc^−/Y^* mice. We observed that OTC-deficient heterozygous female mice tended to be underweight, exhibited mild hyperammonemia, and produced small litters. Administration of the same single AAV vector containing enAsCas12f-HKRA and *hOTC* cDNA to the heterozygous female mice improved their body weight and reduced their blood ammonia levels (Supplemental Figure 6B, C).

Finally, we evaluated the germline transmission of liver-targeted knock-in genome editing using the single AAV vector harboring enAsCas12f-HKRA. The AAV vector genome was primarily detected in the liver but was also detected at low levels in the ovaries of female mice (Supplemental Figure 7A). However, *hOTC* mRNA was scarcely detected in ovaries (Supplemental Figure 7B), and no knock-in sequence could be detected in the liver of offspring derived from the treated female mice. This indicated that germline transmission did not occur (Supplemental Figure 7C).

## Discussion

In this study, we developed a genome-editing strategy that enables the highly efficient knock-in of therapeutic genes using a single AAV vector. This approach uses a compact Cas12f effector, enAsCas12f-HKRA, which is specifically targeted to the liver. Compared with our previous HDR-based strategy (16), this study achieved more efficient knock-in by using NHEJ. This genome-editing strategy led to phenotypic correction in multiple disease models, including for hemophilia B, PC deficiency, and OTC deficiency. The most notable advantage of this knock-in strategy is that all components—including the Cas effector, sgRNA, and donor template—can be packaged into a single AAV vector. This approach eliminates the need for co-delivery of separate Cas effectors and donor vectors into the same cell. It minimizes infection variability, simplifies vector production and quality control, and allows for lower vector doses. In addition, compared with strategies targeting earlier gene regions such as intron 1 (30, 31), our approach targets the region immediately upstream of the stop codon in intron 13, thereby minimizing disruption to the endogenous *Alb* gene. Consistently, we observed no reduction in plasma albumin levels following genome editing. These features collectively offer significant advantages for clinical applications, including enhanced editing efficiency, improved reproducibility, and reduced risk of adverse effects.

The ultimate goal of genome editing is the precise correction of disease-causing variations. Indeed, a landmark clinical study recently demonstrated the curative potential of personalized base editing. In this study, a patient with carbamoyl-phosphate synthetase 1 deficiency—a urea cycle disorder—was successfully treated using mRNA-based base editing delivered via lipid nanoparticles (32). More versatile approaches, such as prime editing are being developed to expand the range of correctable variations beyond the scope of base editing (33). However, these precision editing technologies are not applicable to large deletions, inversions, or other complex variants. Moreover, insufficient editing efficiency may limit their therapeutic potential, particularly in tissues where only a small proportion of cells are edited. In this context, knock-in genome editing offers a more universal and robust strategy. By integrating the therapeutic gene downstream of a strong endogenous promoter—such as the *Alb* promoter, which drives high levels of expression in hepatocytes—this approach enables consistent transgene expression (30), regardless of the underlying variant type. Furthermore, the donor sequence can be flexibly designed to incorporate engineered gene variants with enhanced functionality, such as the FIX R338L variant or the activated PC variant (PC-2RKR), thereby amplifying therapeutic effects.

Knock-in genome editing therapy offers a compelling curative strategy that may replace liver transplantation in certain neonatal-onset inherited liver diseases. Gene replacement therapies that use AAV vectors or lipid nanoparticle-formulated mRNAs have recently emerged as innovative approaches for treating inherited liver disorders (28, 34). mRNA therapy provides a rapid onset of action and avoids genomic integration, making it particularly useful as a bridging strategy before curative interventions, such as genome editing or liver transplantation. AAV-mediated gene replacement therapy enables long-term transgene expression in adults, but its utility in severe neonatal-onset forms is limited because the vector genome becomes diluted as a result of hepatocyte proliferation (35). Although liver transplantation remains a definitive curative therapy for inherited liver diseases, its clinical application is limited by the need for suitable donors and the invasiveness of major surgery. The knock-in genome editing strategy, therefore, has much potential as a standalone alternative and as a complementary option for liver transplantation, particularly for treatment strategies tailored to disease type, severity, and timing of onset.

Several technical and biological challenges must be addressed before this technology can be translated into clinical applications. Although knock-in via NHEJ is cell cycle-independent and generally more efficient than HDR (36), precise control over the orientation and structure of the inserted sequence remains difficult. While the homology-independent targeted integration strategy allows directional insertion of donor sequences (37), its application is limited in our system because of the cleavage characteristics of AsCas12f (38). Reproducibility and sequence fidelity at the target site also require careful validation. For example, co-integration of AAV vector-derived inverted terminal repeat sequences along with the transgene has been reported (39), raising concerns about unintended sequence insertions and structural heterogeneity among cells. Furthermore, optimizing target loci and vector delivery methods may enable this platform to be applied to other organs. Given advances in AAV capsid engineering to enhance organ tropism and the availability of tissue-specific promoters, this platform holds promise for organ-targeted knock-in therapies beyond the liver. Accordingly, extending its application to the treatment of congenital diseases affecting other organs is an important direction for future research, along with the development of scalable, disease-specific strategies.

While this study highlights the therapeutic potential of knock-in genome editing, several limitations warrant further discussion. First, comprehensive analysis of the donor template knock-in patterns was challenging. Although expression of *hF9* and *hOTC* cDNAs in liver hepatocytes was quantified by immunofluorescence staining, multiple variations of the desired knock-in event likely occurred. These include partial insertions of donor fragments and insertions in the reverse orientation relative to the intended direction. Such structural heterogeneity is an inherent limitation of the NHEJ-mediated knock-in approach and comprehensive genome- and transcriptome-wide analyses are necessary to assess the accuracy and uniformity of knock-in events. Second, off-target effects of genome editing could not be evaluated. Sequence differences between human and mouse genomes mean that off-target assessments conducted in mice cannot reliably predict off-target effects in humans. Direct evaluation of off-target effects *in vivo* is difficult, and current strategies primarily depend on predictive assessments performed *in vitro* or *in silico*. Here, GUIDE-seq analysis revealed off-target double-strand breaks induced by several sgRNAs targeting the *ALB* locus in HEK293 cells (Supplemental Figure 8) (40). For future clinical applications, it will be essential to perform such off-target analyses in human-derived cells, supplemented by *in silico* prediction tools (41). Third, long-term expression of the knocked-in therapeutic gene was evaluated only over a limited timeframe. Extended follow-up studies are required to confirm sustained expression and long-lasting therapeutic efficacy. Finally, to enable clinical translation, comprehensive evaluations of editing efficiency, long-term safety, and immunological responses should be conducted in larger animal models, such as non-human primates.

In conclusion, this study demonstrates that knock-in genome editing using enAsCas12f-HKRA and a donor template delivered via a single AAV vector can correct disease phenotypes in multiple monogenic liver disorders. The compact size of AsCas12f, combined with efficient NHEJ-mediated integration into the *Alb* locus, enables therapeutic protein expression through a simplified delivery system. Moreover, this strategy was effective in both neonatal and adult mouse models, without evidence of germline transmission. Overall, the platform developed in this study offers a promising therapeutic approach with the potential to complement or be an alternative to liver transplantation for neonatal-onset inherited liver diseases.

## Methods

### Plasmid construction

cDNAs were inserted into pcDNA4/TO or pcDNA3 (Thermo Fisher Scientific, Waltham, MA) for transient transfection. sgRNAs, driven by the U6 promoter, were incorporated into pBluescript SK-(Agilent Technologies, Inc., Santa Clara, CA) (Supplemental Table 2). *hOTC* cDNAs conjugated with the FLAG sequence were synthesized by AZENTA (Tokyo, Japan), FASMAC (Kanagawa, Japan), and GenScript Japan (Tokyo, Japan). To express an activated form of hPC, a furin-cleaving site was inserted between the light and heavy chain sequences of a *hPROC* cDNA, as described previously (24).

### Cell culture and plasmid transfection

HEK293 cells (JCRB Cell Bank #JCRB9068) and AAVpro 293T cells (Takara Bio, Shiga, Japan) were cultured in Dulbecco’s modified Eagle’s medium (DMEM) (Fujifilm Wako Pure Chemical Corporation, Osaka, Japan) supplemented with 10% fetal bovine serum (FBS; Thermo Fisher Scientific) and GlutaMAX (Thermo Fisher Scientific). Mouse liver-derived TLR-3 cells (JCRB Cell Bank #IFO50380) were cultured in DMEM supplemented with 2% FBS, GlutaMAX, 5 ng/mL human epidermal growth factor (Thermo Fisher Scientific), and Insulin–Transferrin–Ethanolamine supplement (Thermo Fisher Scientific). Cells were transfected with plasmids using Lipofectamine 3000 Reagent (Thermo Fisher Scientific) according to the manufacturer’s instructions.

### T7 endonuclease assay

DNA fragments amplified by PCR using ExTaq DNA polymerase (Takara Bio) were denatured and re-annealed using a thermal cycler, and then treated with T7 endonuclease (Nippon Gene, Tokyo, Japan). The DNA fragments were analyzed and quantified using a microchip electrophoresis system (MCE-202 MultiNA; Shimadzu Corp., Kyoto, Japan).

### AAV vector production

An expression cassette consisting of a promoter, cDNA, and SV40 poly A signal was inserted between the inverted terminal repeat sequences of pAAV. When indicated, enAsCas12f-HKRA, sgRNA driven by the U6 promoter, and donor sequences—including the cDNAs *hF9 (R338L)*, *hPROC*, *hPROC-2RKR*, and *hOTC*— were simultaneously incorporated into an expression cassette in pAAV. To separate the enAsCas12f-HKRA expression cassette from the donor sequence in episomal AAV sequence, an sgRNA recognition sequence was inserted between the expression cassette and the donor sequence (Figure 1A). AAV vectors (serotype 8) were produced using a helper-free system, as described previously (42). Briefly, three plasmids (pHelper [Takara Bio], pRC8 to express Rep and AAV8 capsid, and pAAV) were transfected into AAVpro293T cells (Takara Bio). The AAV vectors were harvested 72 h after the cell transfection, purified by CsCl-based density gradient ultracentrifugation, and quantified by quantitative PCR (qPCR), as previously described (43).

### Animal experiments

C57BL/6J mice were purchased from Japan SLC (Shizuoka, Japan). Hemizygous *Otc^spf-ash^* (sparse fur-abnormal skin and hair) mice and FIX-deficient mice (B6.129P2-*F9^tm1Dws^*) were obtained from The Jackson Laboratory. PC-deficient mice (C57BL/6-*Proc^em1Tsuka^*) were generated as described previously (RIKEN BRC #RBRC11381) (24). *Otc*-deficient C57BL/6 mice lacking exon 2 were generated by genome editing at UNITECH Inc. (Chiba, Japan). Mice were maintained in isolators in the specific pathogen-free facility of Jichi Medical University at 23°C ± 3°C with a 12:12-h light/dark cycle. Mice were fed a standard regular diet (Oriental Yeast Co., Ltd., Tokyo, Japan). When indicated, *Otc^spf-ash^* mice (8–9 weeks of age) were challenged with a high-protein diet (40% kcal) (Oriental Yeast Co., Ltd.). The experimental endpoint for euthanasia under the challenge of the high-protein diet was set as a body weight loss of more than 20% from the initial weight.

Blood was drawn from the jugular vein with a 29G micro-syringe (Terumo Corp., Tokyo, Japan) containing 1/10 (volume/volume) sodium citrate under isoflurane anesthesia. Plasma samples were obtained by centrifugation and stored at −80°C until analysis. For blood ammonia measurement, whole blood (10 µL) was obtained following tail snipping (0.2 mm) of anesthetized mice. AAV vectors were administered intravenously to adult anesthetized mice through the jugular vein (100 µL) or intraperitoneally to neonatal mice (10 µL). For phenotype rescue experiments of *Proc^−/−^*mice by genome editing, an anti-FVIII monoclonal antibody (mAb) (#GMA8015, Green Mountain Antibodies, Burlington, VT) was administered intraperitoneally to pregnant female mice (20 µg/mouse) and to neonatal mice (1 µg/mouse, soon after birth).

### Sex as a biological variable

Sex differences are known to alter the efficacy of gene transduction with the AAV vector in mice (44). Therefore, all experiments featuring AAV vector injection into adult mice were conducted with male mice. However, as the sex of neonatal mice was unknown at birth, all received AAV vectors and were included in the study.

### Quantification of AAV vector genome and mRNA

Genomic DNA was extracted using a DNeasy Blood & Tissue Kit (Qiagen, Hilden, Germany) or GENE PREP STAR (Kurabo Industries Ltd., Osaka, Japan). RNA was extracted using an RNeasy Mini Kit (Qiagen). cDNA was synthesized using a PrimeScript RT Reagent Kit (Takara Bio). qPCR was performed using THUNDERBIRD™ Probe qPCR Mix (TOYOBO, Osaka, Japan) on the QuantStudio™ 12K Flex Real-Time PCR system (Thermo Fisher Scientific). The primers and probes are described in Supplemental Table 3.

### Blood chemistry and coagulation test

Plasma coagulation factor activities were measured using a one-stage clotting assay (CS-1600; Sysmex, Kobe, Japan). hPC:C was measured using an automated coagulation assay (CS-1600; Sysmex) with Berichrom PROTEIN C (Sysmex). Activated partial thromboplastin time was measured using an automated blood coagulation analyzer (CA-500; Sysmex). Plasma albumin concentrations were measured using the Bromocresol Green method with an automated biochemical analyzer by Oriental Yeast Co., Ltd. Blood ammonia was measured using FUJI DRI-CHEM SLIDE NH3-WII (Fujifilm) and a DRI-Chem NX10N analyzer (Fujifilm).

### Measurement of hOTC enzyme activity

Specimens were homogenized in buffer (10 mM HEPES, 0.5% Triton X-100, 2 mM EDTA, 0.5 mM DTT [pH = 7.2]; 100 mg tissue/mL) using a bead homogenizer. After centrifugation, protein concentrations were determined using a protein assay kit (Bio-Rad Laboratories, Inc., Hercules, CA). Liver lysate (10 µg) was incubated with 700 µL reaction mixture (5 mM ornithine, 15 mM carbamyl phosphate, and 270 mM triethanolamine [pH 7.7]) at 37°C for 30 min. The reaction was then stopped by adding 375 µL phosphoric/sulfuric acid solution (3:1) followed by 47 µL 3% 2,3-butanedione monoxime and incubated at 95°C for 15 min. Citrulline production was determined by measuring the absorbance at 490 nm using a Multiskan sky high spectrophotometer (Thermo Fisher Scientific). Results are expressed as mmol citrulline per mg total protein.

### Immunohistochemical analysis

Immunofluorescence analysis was performed as described previously (16). Briefly, sections were blocked in phosphate-buffered saline (PBS) containing 0.3% Triton X-100 and 2% goat serum for 1 h at room temperature, then incubated overnight at 4°C with primary antibodies (anti-hFIX rabbit polyclonal antibody conjugated with biotin [Affinity Biologicals Inc., Ancaster, ON, Canada], anti-DYKDDDDK Tag rabbit mAb [Cell Signaling Technology, Inc., Danvers, MA], and anti-mouse CD146 mAb [BioLegend, Inc., San Diego, CA]). After extensive washing with PBS, sections were incubated for 1 h with Alexa Fluor™ 594 Conjugated Streptavidin (Thermo Fisher Scientific) or species-specific secondary antibodies (anti-rat IgG Alexa Fluor 488 and anti-rabbit IgG Alexa Fluor 594, Thermo Fisher Scientific). Sections were washed again with PBS, mounted with VECTASHIELD Mounting Medium containing DAPI (Vector Laboratories, Newark, CA), and covered with a coverslip. Images were acquired and quantified using a BZ-X700 all-in-one fluorescence microscope (Keyence Corporation, Osaka, Japan).

### Immunoblotting

Cells were lysed in cell lysis buffer (50 mM Tris-HCl (pH 7.5), 150 mM NaCl, 1% deoxycholate, 0.1% sodium dodecyl sulfate, 1% Triton X-100) containing proteinase inhibitor cocktail. The lysate was centrifuged at 20,000 × *g* for 10 min, and the supernatant was recovered. Total protein concentrations were determined using the DC™ Protein Assay Kit (Bio-Rad Laboratories, Inc.). Cell lysates were resolved by SDS polyacrylamide gel electrophoresis and transferred to polyvinylidene fluoride membranes. The membranes were blocked with 5% (weight/volume) skim milk in 20 mM Tris (pH 7.5), 150 mM NaCl, and 0.05% Tween 20 (TBS-T buffer) for 1 h. After extensive washing with TBS-T, the membranes were incubated overnight at 4°C with a mouse anti-FLAG M2 mAb (Sigma-Aldrich, St. Louis, MO) or a β-actin mAb (Santa Cruz Biotechnology, Dallas, TX) in TBS-T containing 5% (weight/volume) bovine serum albumin. Antibody binding was detected using horseradish peroxidase (HRP)-conjugated goat anti-mouse IgG (Bio-Rad) and visualized using Immobilon Western Chemiluminescent HRP Substrate (Millipore, Burlington, MA) and an ImageQuant LAS4000 digital imaging system (GE Healthcare, Buckinghamshire, UK). The bands in the captured images were quantified using ImageJ.

hOTC levels in mouse liver were quantified using a Jess Simple Western System (ProteinSimple, San Jose, CA). Briefly, total liver protein lysates (200 ng/µL) were loaded on the Jess system. hOTC levels were detected using an anti-FLAG M2 antibody conjugated with HRP (Sigma-Aldrich). Lysates of HEK293 cells transfected with the plasmid-expressing hOTC were used as a positive control. hOTC expression levels were standardized using the protein standardization module (ProteinSimple).

### Statistics

The number of independent experiments, biological samples, and mice used are detailed in the figure legends. Statistical analysis was performed using GraphPad Prism 10 software (GraphPad Software Inc., San Diego, CA). The means of two groups were compared using the two-tailed Student’s *t*-test. Multiple comparisons were performed with Tukey’s multiple comparison test. Data are presented as the mean ± standard deviation (SD). Statistical significance was accepted with *p* < 0.05.

### Study approval

All animal experimental procedures were approved by the Institutional Animal Care and Concern Committee of Jichi Medical University (permission number: 23063-03; 23067-01), and animals were cared for in accordance with the guidelines of the committee and ARRIVE.

## Data availability

The data presented in this study are available from the corresponding author upon reasonable request.

## Author contributions

K.B., T.T., and T.O.: designing research studies, conducting experiments, acquiring data, analyzing data, writing the manuscript; Y.K. and N.B.: conducting experiments, acquiring data, analyzing data; K.T., T.S., M.H., K.T., and K.M.: analyzing data; A.H. and O.N.: providing critical reagents. All authors approved the submitted manuscript and agreed to be personally accountable for their own contributions and for the accuracy and integrity of any part of the work.

## Acknowledgements

The study was supported by grants JP25bm1223004, JP25bm1323001, and JP25fk0410061 from the Japan Agency for Medical Research and Development (AMED). We thank Hiromi Ozaki, Yuiko Ogiwara, Sachiyo Kamimura, Yaeko Suto, Mika Kishimoto, Tamaki Aoki, Mai Hayashi, Ayano Suto, Tomoko Noguchi, and Hiroko Hayakawa (Jichi Medical University) for their technical assistance. Some figures were created with Biorender.com. We thank Jeremy Allen, PhD, from Edanz (https://jp.edanz.com/ac) for editing a draft of this manuscript.

## Notes

**Conflicts of interest:** K.B., T.T., N.B., Y.K., M.H., and T.O. are the holders of the patent for the methodology described in the present study. A.H., O.N., and T.O. are the holders of the patent for AsCas12f. No conflicts of interest exist for the other authors.

### Competing Interest Statement

K.B., T.T., N.B., Y.K., M.H., and T.O. are the holders of the patent for the methodology described in the present study. A.H., O.N., and T.O. are the holders of the patent for AsCas12f. No conflicts of interest exist for the other authors.

