## Supplemental materials for "Therapeutic Knock-in Genome Editing Using Single AAV Vectors in Mouse Models of Inherited Liver Disease"

*These authors contributed equally.

**Supplemental Table 1 Codon-optimized human *OTC*-FLAG sequences**
**Supplemental Table 2 Single guide RNAs used in this study**
**Supplemental Table 3 Oligonucleotide primer pairs used in this study**
**Supplemental Figure 1 Efficiency of genome editing targeting the *Alb* locus in TLR-3 cells**
**Supplemental Figure 2 Plasma albumin concentrations in hemophilia B mice treated with a single AAV vector for knock-in genome editing at the *Alb* locus**
**Supplemental Figure 3 Knock-in patterns of cDNA into the *Alb* locus of liver cells via HDR or NHEJ.**
**Supplemental Figure 4 Comparison of anticoagulant function between human PC and PC-2RKR in wild-type adult mice**
**Supplemental Figure 5 Selection of the codon-optimized human *OTC* cDNA**
**Supplemental Figure 6 Rescue of the female *Otc*$^{+/-}$ mouse phenotype by knock-in genome editing with a single AAV vector**
**Supplemental Figure 7 Assessment of germline transmission of knock-in genome editing using a single AAV vector**
**Supplemental Figure 8 Single guide RNAs targeting human *ALB* intron 13**

**Supplemental Table 1. Codon-optimized human *OTC*-FLAG sequences**

| | cDNA sequence |
|---|---|
| CO1 | ATGCTCTTCAACCTGAGGATCTTACTCAACAATGCAGCCTTTAGAAATGGGCACAACTTCATGGTGAGGAA<br>CTTTAGGTGTGGCCAGCCTCTTCAGAATAAAGTGCAGCTGAAAGGCAGAGACTTGCTGACCTTGAAGAAC<br>TTCACTGGAGAGGAAATTAAATACATGCTGTGGCTGTCTGCTGACCTGAAGTTCAGGATAAAGCAGAAAG<br>GTGAATATCTGCCCTTGTTGCAGGGCAAAAGTCTGGGCATGATTTTTGAAAAGAGGAGCACAAGAACCAG<br>GTTGTCCACAGAAACAGGCTTTGCCCTGCTAGGTGGTCATCCCTGCTTCCTTACCACCCAGGACATCCATT<br>TGGGTGTGAATGAAAGTTTAACAGACACAGCCAGAGTTCTCTCTTCAATGGCTGATGCTGTACTGGCCAGG<br>GTCTACAAACAAAGTGACCTGGACACTTTGGCCAAGGAAGCTTCCATCCCAATTATAAATGGCCTCTCAG<br>ATCTCTACCACCCCATCCAGATTTTGGCAGATTATCTCACCCTGCAGGAGCACTACAGTTCTCTAAAAGGG<br>CTTACTTTATCCTGGATTGGAGATGGAAACAACATCCTCCACAGCATTATGATGTCTGCAGCTAAATTTGG<br>AATGCACCTTCAAGCTGCAACACCTAAGGGCTATGAGCCTGATGCATCAGTAACTAAACTTGCTGAGCAG<br>TATGCCAAAGAAAATGGTACAAAGCTGCTGCTCACCAATGATCCCCTGGAGGCTGCCCATGGGGGAAATG<br>TGCTCATCACAGATACCTGGATCAGCATGGGACAAGAAGAGGAGAAGAAGAAAAGACTGCAGGCCTTCC<br>AGGGATACCAGGTGACCATGAAGACTGCAAAAGTTGCAGCCTCAGACTGGACTTTCCTGCACTGTCTGCC<br>AAGGAAGCCTGAGGAGGTGGATGATGAAGTTTTTTATTCCCCCAGAAGCCTGGTGTTTCCTGAAGCAGAA<br>AACAGAAAGTGGACCATCATGGCAGTGATGGTCAGCCTCTTGACTGACTACAGCCCACAGCTCCAAAAGC<br>CAAAGTTTGATTACAAGGACCATGATGGGGACTATAAAGATCATGACATTGATTATAAGGATGATGATGA<br>CAAGTGA |
| CO2 | ATGCTCTTCAACTTGAGAATTCTATTGAATAATGCAGCCTTCAGGAATGGGCATAATTTCATGGTCAGAAA<br>CTTCAGGTGTGGGCAGCCTCTGCAGAACAAAGTTCAACTGAAAGGGAGAGACCTGCTCACATTGAAGAAT<br>TTTACAGGGGAAGAGATTAAATATATGCTGTGGTTATCTGCTGATCTGAAGTTCAGGATAAAACAAAAGG<br>GAGAATACCTCCCCTTGTTACAGGGAAAGTCTCTGGGGATGATATTTGAAAAGAGGTCCACCAGGACAAG<br>ATTGAGTACAGAGACAGGGTTTGCACTCTTGGGTGGCCATCCCTGCTTTCTAACTACCCAGGATATTCATC<br>TGGGAGTCAATGAGAGTCTTACTGATACTGCAAGGGTCTTAAGTTCAATGGCTGATGCTGTCCTTGCAAGA<br>GTGTACAAACAATCAGACCTGGATACCCTTGCTAAAGAGGCATCAATTCCAATCATCAATGGACTTTCTGA<br>TCTATACCACCCCATCCAGATCCTTGCTGACTACCTTACACTCCAAGAACATTACTCAAGCCTCAAAGGAC<br>TTACTCTGTCCTGGATTGGGGATGGCAATAACATTCTCCACTCCATCATGATGAGTGCAGCTAAGTTTGGC<br>ATGCACCTCCAGGCTGCCACCCCTAAGGGTTATGAGCCAGATGCTTCTGTTACCAAACTGGCTGAGCAGTA<br>TGCTAAAGAAAATGGCACCAAGCTGCTGCTGACTAATGACCCACTGGAGGCAGCCCATGGTGGCAATGTA<br>CTGATTACAGATACATGGATCAGCATGGGACAGGAGGAGGAAAAGAAGAAAAGGCTGCAGGCCTTTCAG<br>GGATACCAGGTTACCATGAAGACAGCCAAAGTGGCTGCTAGTGACTGGACCTTTCTGCACTGCCTGCCTA<br>GGAAGCCTGAGGAGGTGGATGATGAAGTGTTCTATAGCCCCAGGAGCCTGGTGTTTCCTGAGGCTGAAAA<br>CAGGAAATGGACTATCATGGCAGTGATGGTGTCTCTGCTGACAGATTATTCTCCTCAGCTGCAGAAGCCCA<br>AGTTTGACTACAAGGACCATGATGGAGACTACAAGGACCATGACATTGACTATAAAGATGATGATGACAA<br>GTGA |
| CO3 | ATGCTGTTCAACCTGAGAATCCTGCTGAACAATGCAGCCTTCAGAAATGGCCACAACTTCATGGTGAGAA<br>ACTTCAGATGTGGCCAGCCTCTGCAGAACAAAGTGCAACTGAAGGGCAGAGACCTGCTGACCCTGAAGAA<br>CTTCACAGGAGAAGAGATCAAGTACATGCTGTGGCTGTCAGCAGACCTGAAGTTCAGAATCAAACAGAAG<br>GGAGAGTACCTGCCTCTGCTGCAGGGCAAGAGCCTGGGCATGATCTTTGAGAAGAGGAGCACCAGAACCA<br>GACTGTCCACAGAAACTGGCTTTGCCCTGCTGGGTGGCCACCCCTGCTTCCTCACCACCCAGGACATCCAC<br>CTGGGAGTGAATGAGAGCCTGACAGACACAGCCAGGGTGTTAAGCAGCATGGCTGATGCTGTGCTGGCCA<br>GAGTGTACAAGCAGTCTGACCTGGACACACTGGCCAAGGAGGCCAGCATCCCCATCATCAATGGCCTGTC<br>TGACCTGTACCACCCAATCCAGATCCTGGCTGACTACCTGACCCTGCAGGAGCACTACAGCTCCCTGAAGG<br>GCCTGACCCTGAGCTGGATAGGAGATGGCAACAACATCCTGCACAGCATCATGATGTCTGCTGCCAAGTT<br>TGGCATGCACCTGCAGGCTGCCACCCCTAAGGGCTATGAACCTGATGCCTCTGTTACCAAGCTGGCAGAA<br>CAGTATGCCAAGGAGAATGGCACCAAACTGCTACTGACCAATGACCCCCTGGAAGCTGCCCATGGAGGCA<br>ATGTCCTGATCACAGACACCTGGATCAGCATGGGCCAGGAGGAAGAGAAGAAAAAAAGACTGCAGGCCT<br>TCCAAGGCTACCAGGTGACCATGAAGACAGCCAAAGTGGCAGCCAGTGACTGGACCTTCCTCCACTGCCT<br>GCCTAGAAAGCCTGAAGAGGTGGATGATGAGGTGTTCTACAGCCCTAGAAGCCTGGTGTTCCCAGAGGCT<br>GAAAACAGGAAGTGGACCATCATGGCTGTGATGGTGAGCCTGCTGACAGACTACAGCCCTCAACTGCAGA<br>AGCCCAAGTTTGACTACAAGGACCATGATGGGGACTACAAGGACCATGACATTGACTACAAGGATGATGA<br>TGACAAGTGA |

**Supplemental Table 2. Single guide RNAs used in this study**

| Target gene | sgRNA | Sequence | PAM sequence |
|---|---|---|---|
| *Alb* intron 13 | sgRNA1 | GTTTCAAATTTGTGACACAG | TTTG |
| | sgRNA2 | TGACACAGAAGAGCATAGTT | TTTG |
| | sgRNA3 | GGCACAACAGATGTCAGAGA | TTTG |
| | sgRNA4 | CACTTCATAGCCATAGGCAA | TTTG |
| | sgRNA5 | AAACCAAATAGTGATAATAG | TTTG |
| | sgRNA6 | CAAGTATTTCTAACTATGCT | TTTG |
| | sgRNA7 | GACCCTGAAAACAAAGAAAT | TTTG |
| *Alb* intron 14 | sgRNA1 | ACAAGAGCCATACAGACAAG | TTTG |
| | sgRNA2 | TAGATAAGAAACTGAACATA | TTTG |
| | sgRNA3 | TGTAAATTGCATTAACCTAG | TTTG |
| *ALB* intron 13 | sgRNA1 | GGGACAACTATGTCCGTGAG | TTTG |
| | sgRNA2 | GTTAGGCTAGGGCTTAGGGA | TTTG |
| | sgRNA3 | TACATGTGGGACAGGGATCT | TTTG |
| | sgRNA4 | CATGTTTGGTTAGGCTAGGG | TTTG |
| | sgRNA5 | ATATATAAATCCCTAAGCCC | TTTG |
| | sgRNA6 | TAAAATAAGATCCCTGTCCC | TTTG |
| | sgRNA7 | TGATGCTTATGAATATTAAT | TTTG |
| *Otc intron 1* | sgRNA1 | ACTTGTATGCCTTTTGTCAG | TGG |
| *Otc intron 2* | sgRNA2 | TAAGTGCAGTGCATAGCCAT | GGG |

**Supplemental Table 3. Oligonucleotide primer pairs used in this study**

| Target gene | | Sequence |
|---|---|---|
| T7 endonuclease assay | | |
| *Alb* intron 13 | F | 5'-CCTGCTTCTCGACTGAGGTC-3' |
| | R | 5'-CCCTGGTTTGGTCTCCTTTT-3' |
| *Alb* intron 14 | F | 5'-GCCACACTGCTGCCTATTAAATACC-3' |
| | R | 5'-GGACTCCACATAGTGGTTCATGTAAG-3' |
| *ALB* intron 13 | F | 5'-TGCACTTGTTGAGCTCGTGAAACAC-3' |
| | R | 5'-CTGAGATGCTTTTAAATGTGATGTTATAAGCCTAAGGCAGCTT-3' |
| Real time qPCR | | |
| SV40 poly A | F | 5'-AGCAATAGCATCACAAATTTCACAA-3' |
| | R | 5'-CCAGACATGATAAGATACATTGATGAGTT-3' |
| | Probe | 5'-AGCATTTTTTTCACTGCATTCTAGTTGTGGTTTGTC-3' |
| *OTC* mRNA | F | 5'-GGATAGGAGATGGCAACAACA-3' |
| | R | 5'-AACAGAGGCATCAGGTTCATAG-3' |
| | Probe | 5'-ATGATGTCTGCTGCCAAGTTTGGC-3' |
| *Gapdh* mRNA | | Mm99999915_g1 (Thermo Fisher Scientific) |
| Detection of HDR and NHEJ | | |
| *Alb- human F9* mRNA | F | 5'-CTCAGGTGTCAACCCCAACT-3' |
| | R | 5'-TACCTCTTTGGCCGATTCAG-3' |
| *Gapdh* mRNA | F | 5'-CCATCACCATCTTCCAGGAG-3' |
| | R | 5'-CCTGCTTCACCACCTTCTTG-3' |
| human *F9* knock into *Alb* intron 13 (NHEJ) | F | 5'-CGTCATGGGTGTGACTTTTG-3' |
| | R | 5'-TACCTCTTTGGCCGATTCAG-3' |
| human *F9* knock into *Alb* intron 14 (HDR) | F | 5'-CGTCATGGGTGTGACTTTTG-3' |
| | R | 5'-TACCTCTTTGGCCGATTCAG-3' |
| mouse *F8* exon 18 and | F | 5'-TGGCAGTGTACAACCTCTACC-3' |
| intron 18 (Internal control) | R | 5'-AAGGGACAAATGGAGAACTGA-3' |
| *OTC* knock into *Alb* | F | 5'- GCTACAGCGGAGCAACTGAAGACTGTCATG-3' |
| intron 13 (NHEJ) | R | 5'- AACAGAGGCATCAGGTTCATAG-3' |

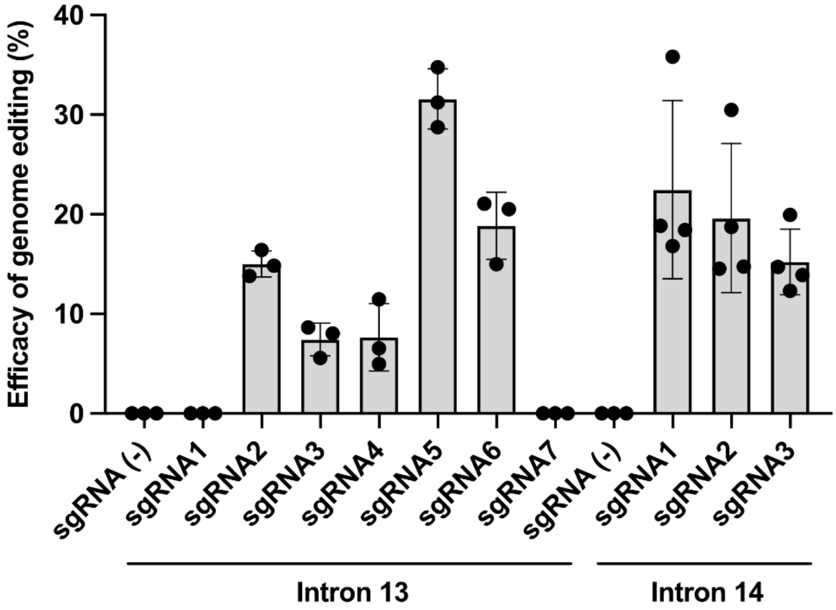


**Supplemental Figure 1. Efficiency of genome editing targeting the *Alb* locus in TLR-3 cells.** TLR-3 cells were transfected with plasmids expressing enAsCas12f-HKRA and single guide RNA (sgRNA) [sgRNA(-); enAsCas12f-HKRA only] targeting *Alb* intron 13 or intron 14. DNA double-strand breaks induced by enAsCas12f-HKRA were evaluated by T7 endonuclease assay. Values represent the mean ± SD (n = 3–4). $P = 0.2620$ was determined by one-way ANOVA with Tukey's multiple comparisons test compared with sgRNA5 (intron 13) and sgRNA1 (intron 14).

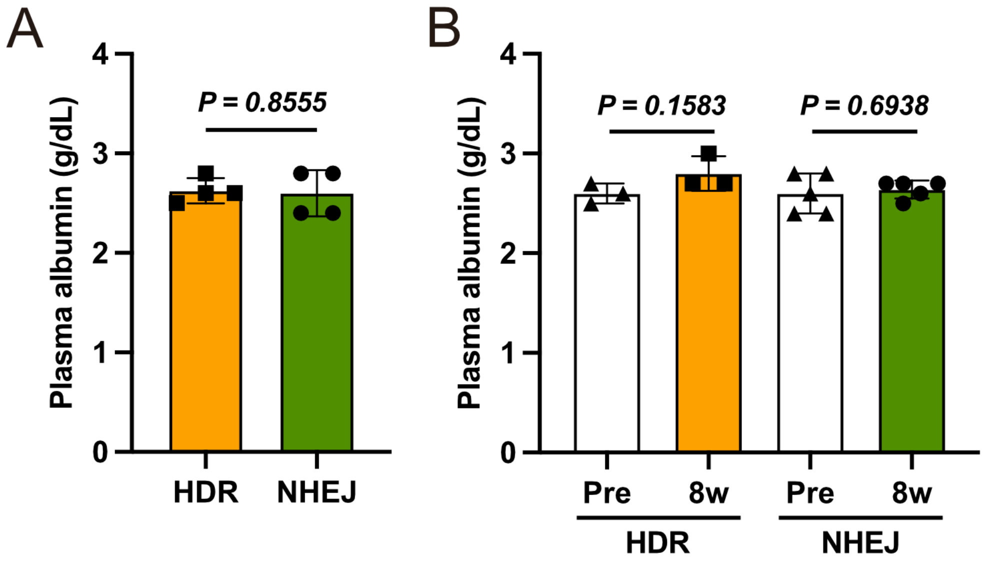


**Supplemental Figure 2. Plasma albumin concentrations in hemophilia B mice treated with a single AAV vector for knock-in genome editing at the *Alb* locus.** (**A**) Neonatal hemophilia B mice were treated with the single AAV vector for knock-in genome editing of *hF9* R338L cDNA into the *Alb* locus via double-strand breaks at intron 13 (NHEJ) or intron 14 (HDR) ($3 \times 10^{11}$ vg/mouse). Plasma albumin concentrations were measured at 8 weeks after vector injection. The values are the mean ± SD (n = 4). (**B**) Adult hemophilia B mice were treated with the single AAV vector for knock-in genome editing of *hF9* R338L cDNA into the *Alb* locus via double-strand breaks at intron 13 (NHEJ) or intron 14 (HDR) ($1 \times 10^{12}$ vg/mouse). Plasma albumin concentrations were measured at 0 (pre) and 8 weeks (8 w) after vector injection. The values are the mean ± SD (n = 3–5). The *P* values between two groups were analyzed by the two-tailed Student's *t*-test. HDR, homology-directed repair; NHEJ, non-homologous end joining.

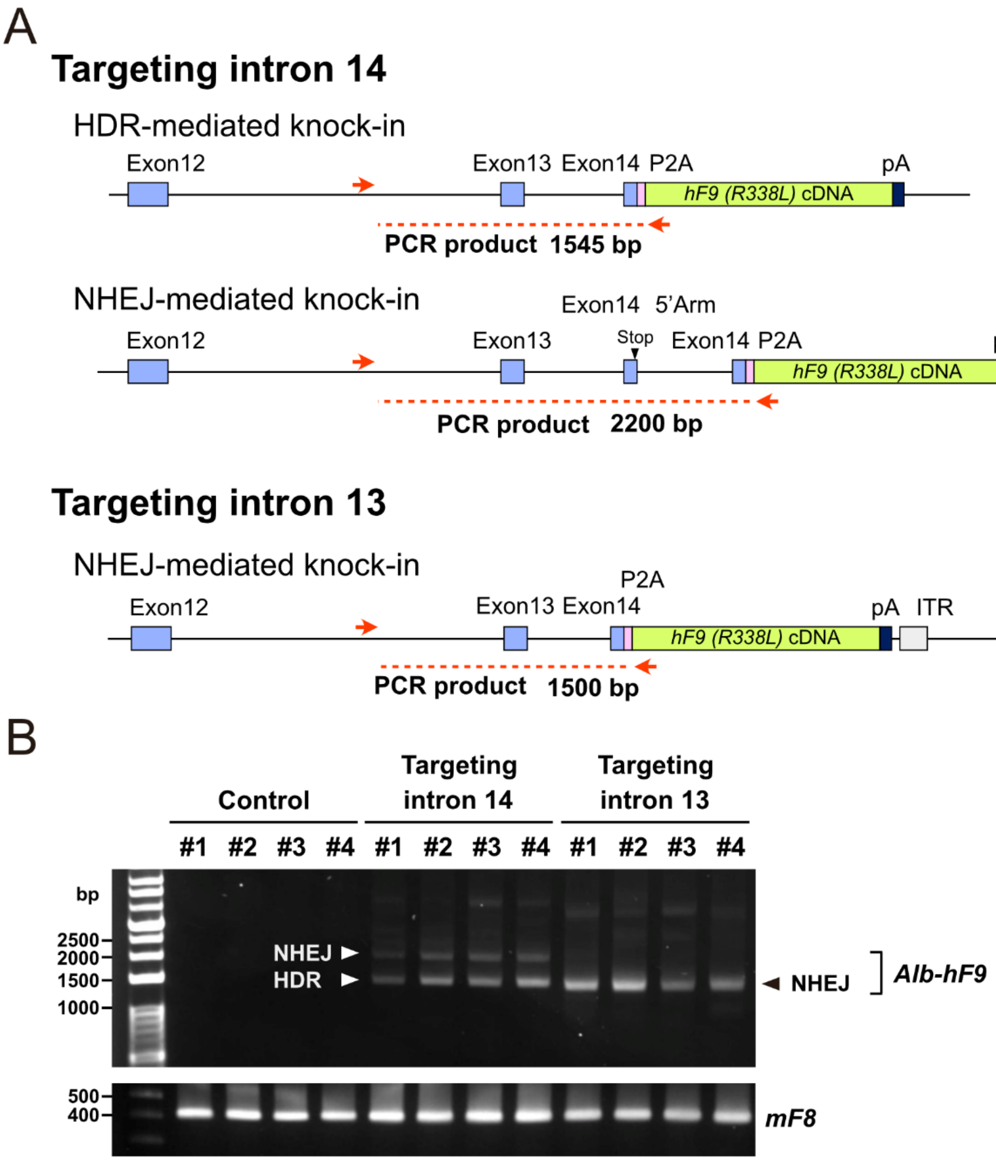


**Supplemental Figure 3. Knock-in patterns of cDNA into the *Alb* locus of liver cells via HDR or NHEJ.** Neonatal hemophilia B mice were treated with the single AAV vector for knock-in genome editing of *hF9* R338L cDNA into the *Alb* locus via double-strand breaks at intron 13 (NHEJ) or intron 14 (HDR) ($3 \times 10^{11}$ vg/mouse). Liver DNA was harvested for PCR analysis 12 weeks after the vector injection. (**A**) Schematic presentation of possible insertion patterns of the donor sequences at the *Alb* locus by targeting intron 14 and intron 13. The arrows indicate PCR primers. (**B**) Insertion of the *hF9* cDNA at the *Alb* locus in liver was identified by PCR. HDR and NHEJ insertion can be distinguished by the size of PCR products. The *mF8* gene was used as a control for genomic DNA amplification. The # numbers indicate individual mice (n = 4). HDR, homology-directed repair; NHEJ, non-homologous end joining; ITR, inverted terminal repeat; *mF8*, mouse coagulation factor VIII gene.

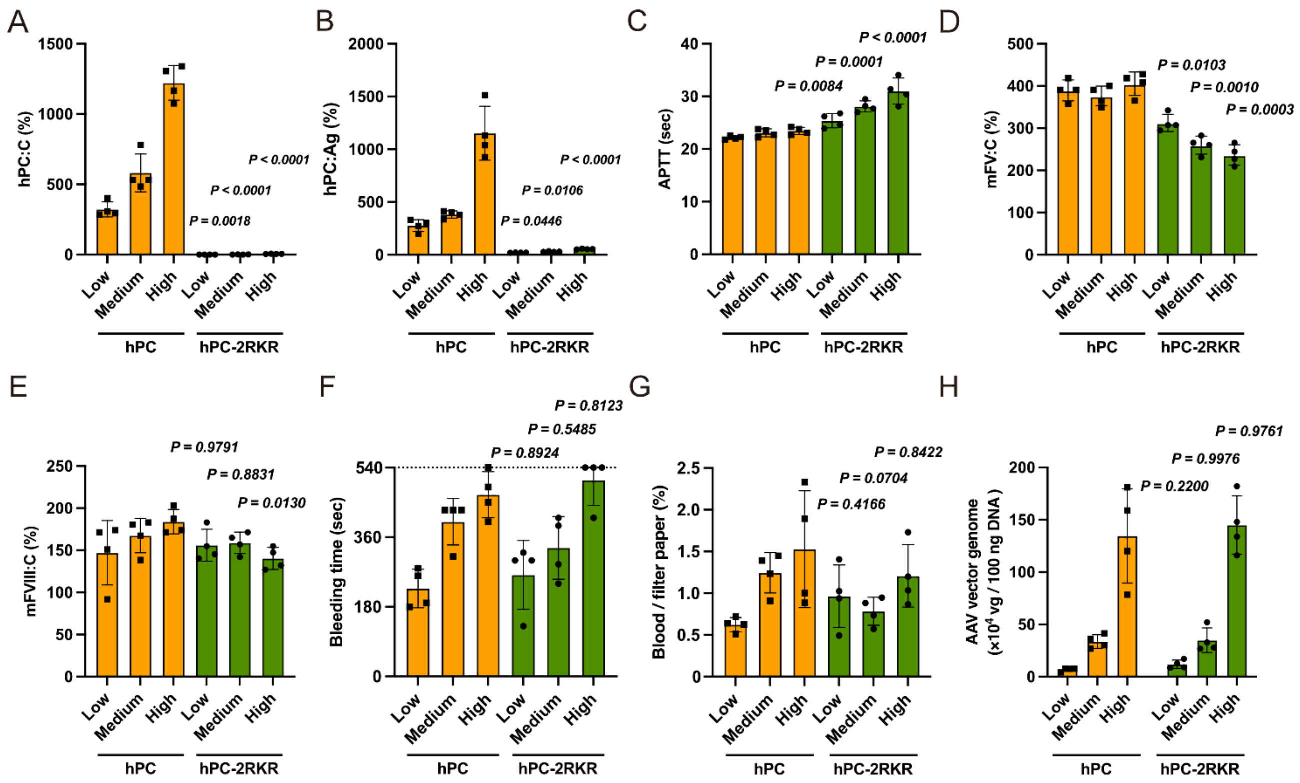


**Supplemental Figure 4. Comparison of anticoagulant function between human PC and PC-2RKR in wild-type adult mice.** Adeno-associated virus (AAV) vectors expressing wild-type human PC (hPC) or the engineered activated hPC (hPC-2RKR) under control of the HCRhAAT promoter were intravenously administered into 7-week-old C57BL/6J male mice at low, medium, and high doses: $4.0 \times 10^{10}$, $1.2 \times 10^{11}$, and $4.0 \times 10^{11}$ vg/mouse, respectively. (**A, B**) Plasma hPC activity (hPC:C) and hPC antigen (hPC:Ag) levels at 4 weeks after vector injection. (**C–E**) Activated partial thromboplastin time (APTT; **C**), mouse factor V activity (mFV:C; **D**), and mouse factor VIII activity (mFVIII:C; **E**) were measured using an automated coagulation analyzer. (**F, G**) Bleeding time and blood volume of tail clip assays. (**H**) AAV vector genome in liver tissue at 6 weeks post-injection was measured using qPCR. Values represent the mean ± SD (n = 4). The *P* values were determined using two-way ANOVA with Šídák's multiple comparisons test comparing hPC and hPC-2RKR at the same vector dose. 2RKR, RKRRKR; HCRhAAT, a chimeric promoter consisting of an enhancer element of the hepatic control region of the *Apo E/C1* genes and the human anti-trypsin promoter.

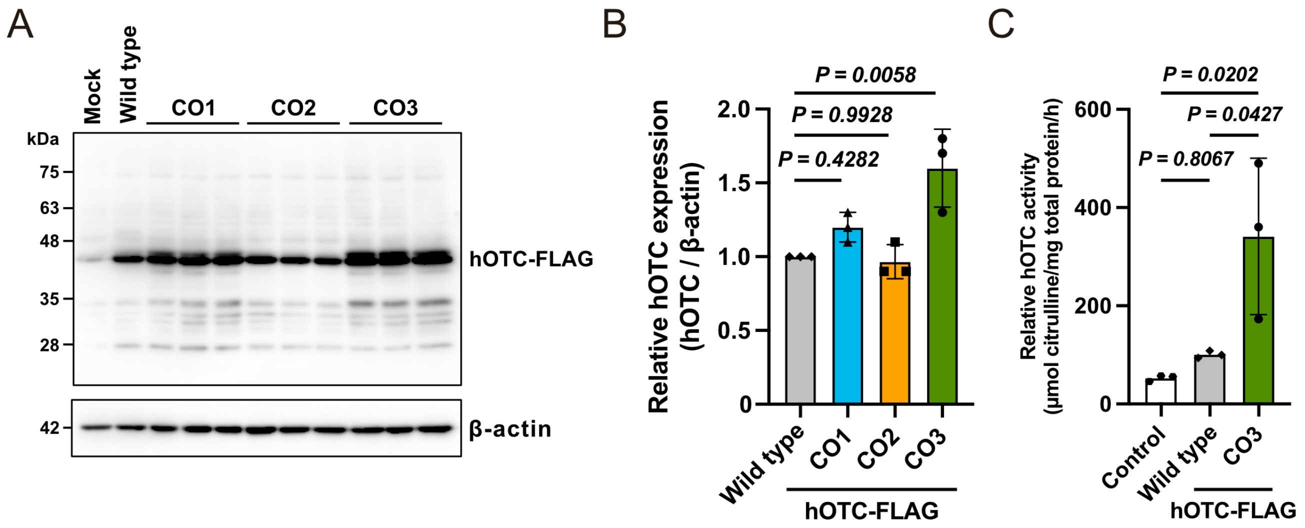


**Supplemental Figure 5. Selection of the codon-optimized human *OTC* cDNA.** HEK293 cells were transfected with pCDNA3 harboring wild-type or codon-optimized (CO) *hOTC-FLAG* cDNA. (**A**) hOTC protein in cell lysates was analyzed by immunoblotting with an anti-FLAG antibody. (**B**) Quantification of hOTC protein levels, normalized against levels of β-actin. Values represent the mean ± SD (n = 3). *P* values were determined using one-way ANOVA and Tukey's multiple comparisons test for comparison with wild-type OTC sequence. (**C**) OTC enzyme activity in transfected HEK293 cell lysates. Control: untransfected HEK293 cells. Values represent the mean ± SD (n = 3). *P* values were determined using one-way ANOVA with Tukey's multiple comparisons test.

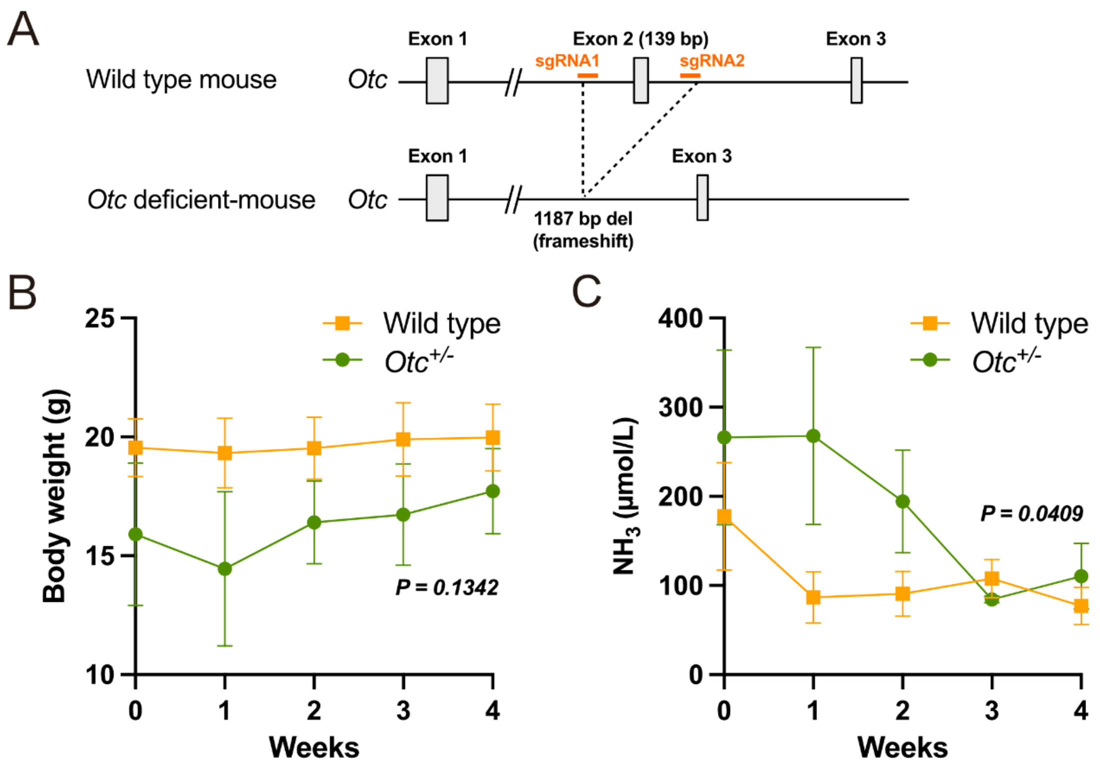


**Supplemental Figure 6. Rescue of the female *Otc*$^{+/-}$ mouse phenotype by knock-in genome editing with a single AAV vector.** (**A**) Generation of *Otc*-deficient mice. Single guide RNAs targeting *Otc* intron 1 and intron 2, together with SpCas9, were introduced into C57BL6/J mouse zygotes by electroporation. (**B**, **C**) Adult *Otc*$^{+/-}$ female mice or litter-matched wild-type females were treated with the single AAV vector for knock-in genome editing to insert an *hOTC* cDNA (CO3) at intron 13 (NHEJ) of the *Alb* locus ($1 \times 10^{12}$ vg/mouse). Changes in body weight (**B**) and blood ammonia ($NH_3$) concentrations (**C**) were assessed after vector injection. Values represent the mean ± SD (n = 4). The *P* values were determined by two-way ANOVA.

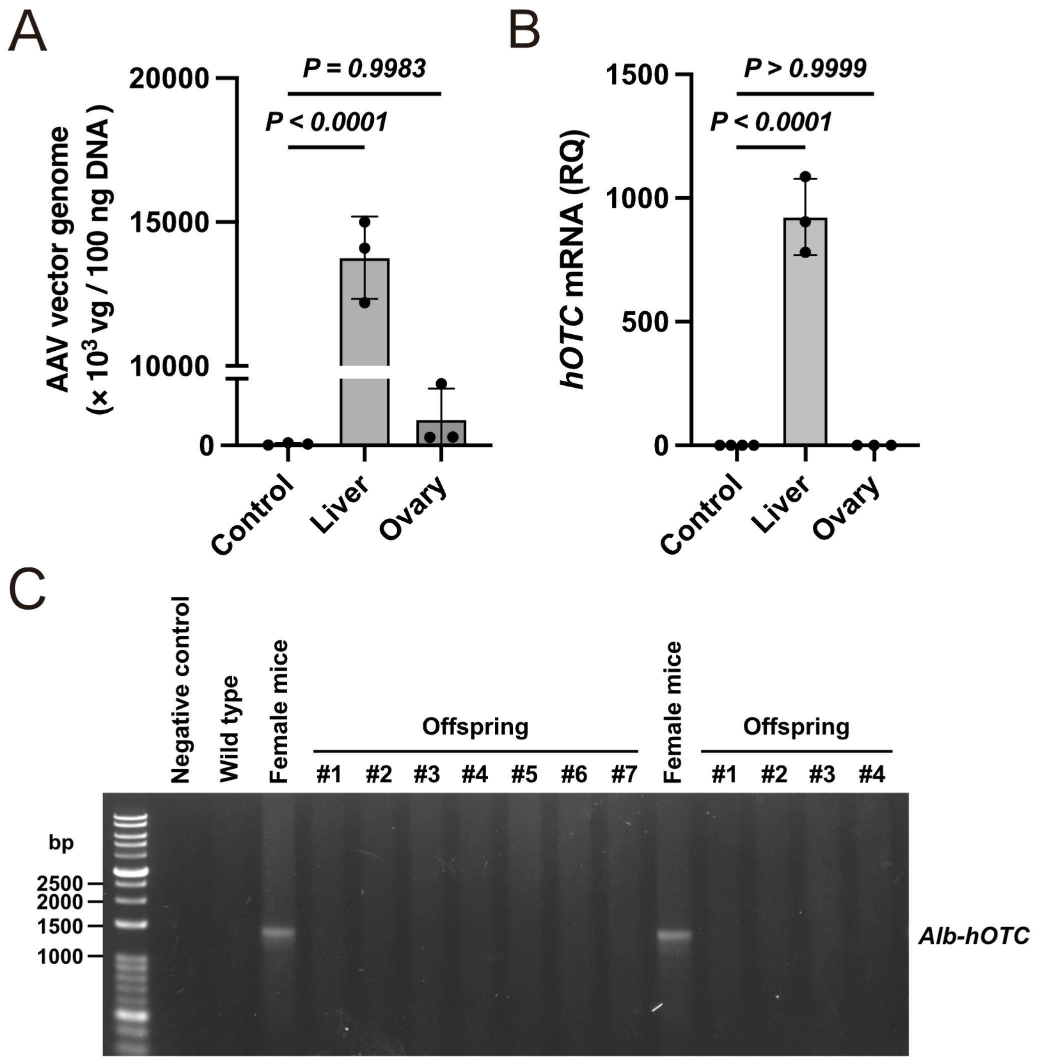


**Supplemental Figure 7. Assessment of germline transmission of knock-in genome editing using a single AAV vector.** (**A, B**) Adult wild-type mice were treated with the single AAV vector for knock-in genome editing to insert an *hOTC* cDNA (CO3) at intron 13 of the *Alb* locus ($1 \times 10^{12}$ vg/mouse). Liver and ovaries were harvested 8 weeks after vector injection. Quantification of AAV vector genome (**A**) and *hOTC* mRNA levels (**B**) in tissues. The *P* values were determined by one-way ANOVA with Tukey's multiple comparisons test compared with control (untreated mice). (**C**) PCR analysis to detect the knock-in sequence in liver DNA obtained from female mice treated with the single AAV vector for knock-in genome editing and from their offspring.

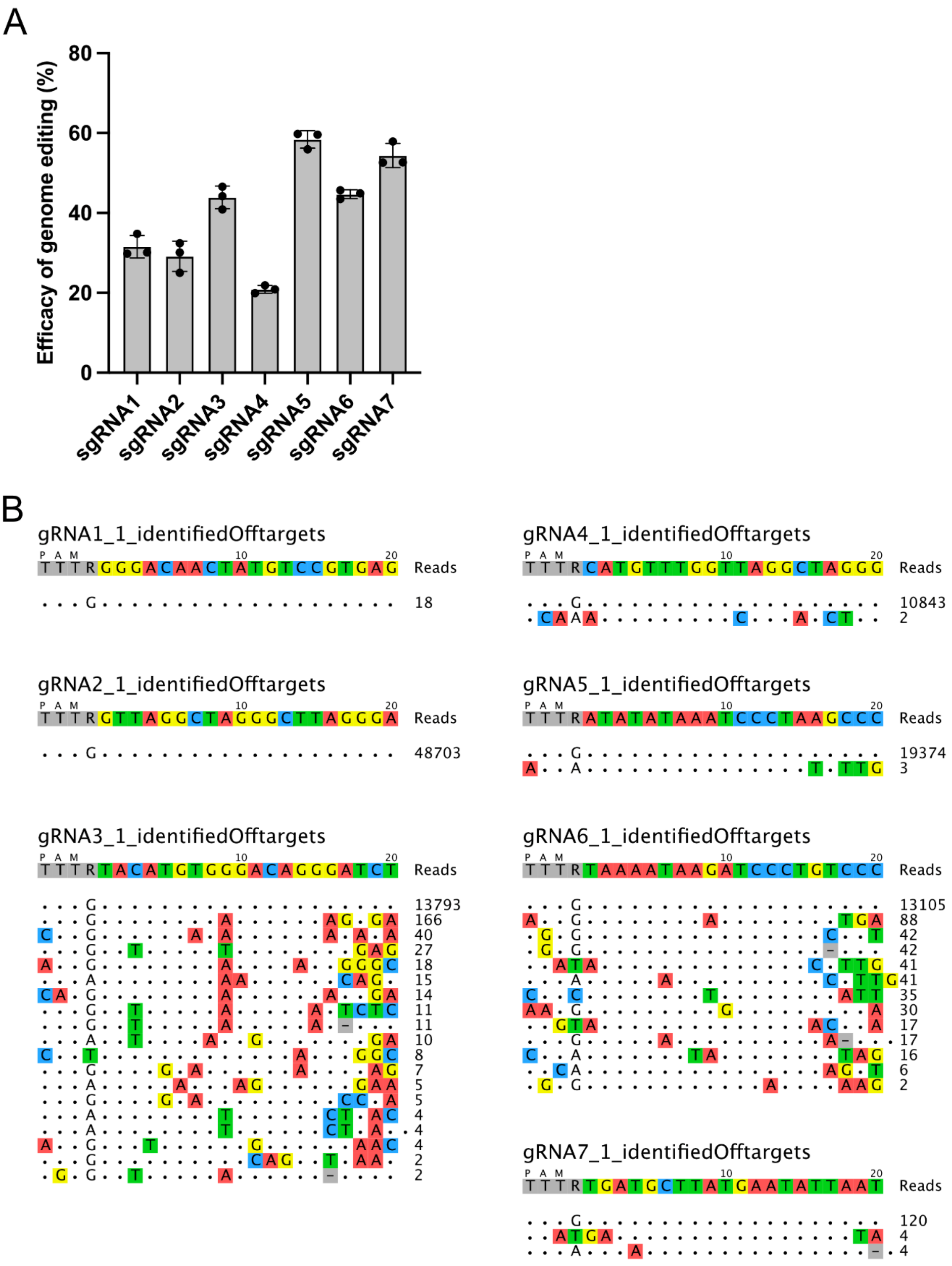


**Supplemental Figure 8. Single guide RNAs targeting human *ALB* intron 13.** (**A**) HEK293 cells were transfected with plasmids expressing enAsCas12f-HKRA and single guide RNA (sgRNA) targeting human *ALB* intron 13. Efficacy of genome editing was evaluated by T7 endonuclease assay. Data are expressed as the mean ± SD (n = 3). (**B**) Off-target sites of each sgRNA were identified by GUIDE-seq analysis.
